# Mating system and speciation I: accumulation of genetic incompatibilities in allopatry

**DOI:** 10.1101/2022.07.25.501356

**Authors:** Lucas Marie-Orleach, Christian Brochmann, Sylvain Glémin

## Abstract

Self-fertilisation is widespread among hermaphroditic species across the tree of life. Selfing has many consequences on the genetic diversity and the evolutionary dynamics of populations, which may in turn affect macroevolutionary processes such as speciation. On the one hand, because selfing increases genetic drift and reduces migration rate among populations, selfing may be expected to promote speciation. On the other hand, because selfing reduces the efficacy of selection, selfing may be expected to hamper ecological speciation. To better understand under which conditions and in which direction selfing affects the build-up of reproductive isolation, an explicit population genetics model is required. Here, we focus on the interplay between genetic drift, selection and genetic linkage by studying speciation without gene flow. We test how fast populations with different rates of selfing accumulate mutations leading to genetic incompatibilities. When speciation requires the population to pass through a fitness valley caused by underdominant and compensatory mutations, selfing reduces the depth and/or breadth of the valley, and thus overall facilitates the fixation of incompatibilities. When speciation does not require the population to pass through a fitness valley, as for Bateson-Dobzhanzky-Muller incompatibilities (BDMi), the lower effective population size and higher genetic linkage in selfing populations facilitates the fixation of incompatibilities. Interestingly, and contrary to intuitive expectations, local selection does not always accelerate the build-up of reproductive isolation in outcrossing relative to selfing populations. Our work helps to clarify how selfing lineages may speciate and diversify over time, and emphasizes the need to account for interactions among segregating mutations within populations to better understand macroevolutionary dynamics.

**Author summary:** Hermaphroditic organisms may use their male gametes to fertilise their own female gametes, and species vary greatly in how much they self-fertilise. Self-fertilisation induces many genetic modifications in the population, which may ultimately affect the rates at which lineages diversify. Here we aim to build predictions on how self-fertilisation affects the rate at which reproductive isolation arises between geographically isolated populations. Specifically, we develop theoretical models in which populations varying in their rates of self-fertilisation may fixate mutations leading to reproductive isolation. We first explored scenarios in which reproductive isolation is made by mutations whose fixations necessitate the population to experience temporally deleterious effects (i.e., a fitness valley), and found that self-fertilisation reduces the breadth and depth of the fitness valley and thereby overall facilitates the accumulation of such mutations. Second, we explored scenarios in which genetic incompatibilities are caused by interactions between derived alleles of different genes (i.e., BDMi). By allowing the BDMi to occur within populations, we found that self-fertilisation reduces the manifestation of BDMi within population, and thereby facilitates their fixation. This effect prevails even in the face of local adaptation. Thus, our study clarifies how fast species are expected to arise in self-fertilisation lineages.

## Introduction

> *Species belonging to the same section or species group usually cross freely in woody plants and perennial herbs, but are usually separated by incompatibility barriers in annual herbs. [*…*] It is possible, therefore, that the correlation [*…*] between sterility and life form is a reflection of a more fundamental relationship between type of breeding system and the formation of sterility barriers”* [1].

The wide variety of mating systems observed in animals and plants, and also fungi and algae, has multiple ecological and evolutionary consequences that might impact higher-level evolutionary processes such as species extinction and speciation. For instance, hermaphroditic species vary in their rate of self-fertilisation – spanning from obligate outcrossing to predominant selfing species, and including all degrees of mixed mating [2–4] – which has long been argued to affect macroevolutionary processes [5–9]. Because selfing tends to reduce both the genetic diversity and the population adaptive potential, selfing lineages have been argued to be ‘evolutionary dead-ends’ as they are expected to go extinct at faster rates than outcrossing lineages [5, 8, 10, 11]. The study of the macroevolutionary effects of selfing has however mostly focused on species extinction, while the effects of selfing on speciation has received relatively less attention, both empirically and conceptually (but see [7, 9]).

The effects of selfing on speciation have been studied based on phylogenies. Phylogenetic trees with variation in mating system may allow us to estimate and compare the rates of species diversification, and potentially speciation and extinction, in selfing *vs*. outcrossing lineages (*e*.*g*., [12–15]). For instance, in the Solanaceae plant family, outcrossing is enforced by a self-incompatibility mechanism that has broken down several times, leading to multiple independent self-compatible lineages. Compared to the self-incompatible lineages (*i*.*e*., obligate outcrossers), the self-compatible ones (*i*.*e*., potential selfers) show lower diversification rates [16] which, interestingly, are likely to be caused by higher rates of both speciation and species extinction in the selfing lineages [12]. Other phylogenetic studies carried out in the Primulaceae [13] and in the Onagraceae [15] plant families suggest that young selfing taxa experience a burst of speciation that fade away with time (*i*.*e*., a ‘senescing diversification rate’ [17]). In contrast, mixed evidence are reported in the Polemoniaceae plant family, in which alternative phylogenetic methods provide positive or no associations between selfing and speciation rates [14].

The effects of selfing on speciation have also to some degree been studied based on experimental crosses, addressing whether reproductive isolation (RI) between populations evolves at different rates in selfing *vs*. outcrossing species. To our knowledge however, there are only a few of such studies. For instance, in the Arctic flora, intraspecific crosses between geographically isolated populations of eight predominantly selfing species resulted in F1 hybrids with low pollen fertility and seed set, whereas no reduced fertility was observed in the single outcrossing species for which successful crosses could be made [18, 19]. In the selfing species, it was estimated that RI may have developed over just a few millennia. The experimental crosses in one of these Arctic species, *Draba nivalis*, were also used to address the genetic architecture of RI. Quantitative trait loci analyses of F2 populations of this predominantly selfing species showed that the post-zygotic incompatibilities are due to single-locus underdominance, a putative chromosomal translocation, and nuclear-nuclear and cyto-nuclear epistatic incompatibilities [20, 21].

Theoretical expectations on the effects of selfing on speciation are poorly studied, and not straightforward. This is because selfing impacts several interconnected population genetics parameters that have opposite effects on speciation. First, selfing decreases gene flow within and among populations, enhancing the isolation of populations and thus possibly facilitating speciation [7]. Second, the non-random sampling of gametes used for reproduction by selfing individuals reduces the effective population size. For instance, the effective population size is expected to be halved in purely selfing species compared to a randomly mating outcrossing species of the same population size [22, 23]. A reduction of effective population size has cascading effects. It elevates genetic drift, reduces genetic polymorphism, and overall weakens selection. Third, selfing increases homozygosity, which makes recombination less efficient because homologous chromosomes tend to be identical [23]. Thus, recombination breaks down linkage disequilibrium less efficiently in strongly selfing populations, thereby reducing the evolutionary advantages of recombination [24], and overexposing the populations to the deleterious effects of linked selection, such as background selection [25], further reducing effective population size in selfing populations [26, 27].

Here, we developed analytical and simulation models of population genetics to better understand the effects of selfing on speciation. We did not consider the effects of gene flow, but focused on the interplay between genetic drift, genetic linkage, and selection efficacy. We studied how mutations leading to RI accumulate within populations differing in selfing rates, and asked if selfing affects (i) the pace of speciation, and (ii) the genetic architecture of RI.

We sequentially explored three types of mutations: underdominant mutations, compensatory mutations, and Bateson-Dobzhansky-Muller incompatibility mutations. Underdominant mutations have deleterious effects in the heterozygous state, but have no deleterious effects in either homozygous state (which may for instance be due to structural variants [28]). Compensatory mutations are a pair of mutations that are both deleterious when they occur alone in a genome, but are neutral when they occur together [29] (*e*.*g*., compensatory evolution of *cis*- and *trans*-regulation of gene expression [30–32]). Finally, Bateson-Dobzhansky-Muller incompatibility (BDMi) mutations are a pair of mutations that have no deleterious effects when they occur alone in a genome, but cause genetic incompatibilities when they occur together [33–36]. Importantly, the fixation of underdominant and compensatory mutations requires the population to pass through a fitness valley in which mutations may be counter selected. In contrast, the fixation of BDMi mutations may be neutral, or even positively selected when the mutations are advantageous [37]. Because homozygotes are formed more readily in selfing species, selfing has previously been shown to facilitate the fixation of underdominant mutations [38]. It is however unknown if and how selfing modulate the accumulation of mutations with epistatic effects, such as compensatory and BDMi mutations.

Overall, we hypothesised that the effects of selfing on speciation depend on the mode of speciation. If RI evolves through genetic drift, selfing should promote speciation because underdominant and compensatory mutations are more likely to get fixed through genetic drift in selfing lineages. In contrast, if RI evolves as a by-product of selection (*e*.*g*., ecological speciation), we expect genetic incompatibilities to arise more readily in outcrossing lineages because selfing populations have an overall lower efficacy of selection.

## Methods

So far, the accumulation of BDMi in allopatry has mostly been modelled as a combinatorial process of substitutions, each predicted by single-locus theory [36, 37]. Extending such results to selfing populations would be straightforward but partly misleading as these models do not explicitly consider the underlying multi-loci population genetics dynamics and the possible interactions among alleles that can be affected by selfing. Instead, we studied the effects of selfing on speciation by modelling – in a single population – the fate of different types of mutation that create genetic incompatibilities among populations. Especially, we determined the probability of and the time to fixation of the different categories of incompatibilities. The fixation of a single incompatibility mutation is, in most cases, not sufficient to complete speciation but determines the overall pace at which RI builds. We first studied the dynamics of a single (pair of) incompatibility and then consider multi-loci models where mutations can occur recurrently throughout the genome.

For all models, we considered a population of *N* hermaphroditic individuals reproducing by selfing at rate 0 ≤ *σ* ≤ 1. The effective size of partially selfing populations is given by Pollak [22]:

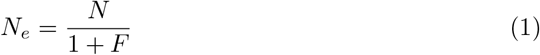

where *F*, the Wright’s fixation index, is:

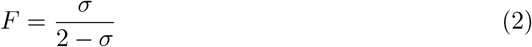

Note that the effective population size, *N*_*e*_, can be further reduced in selfing populations due to background selection, which we included in the single-locus and two-loci simulations (see Simulations). For multi-loci models, we assumed a recombination rate *r* between adjacent loci and an equal mutation rate, *µ*, for all loci. We first analysed the single-locus and two-loci models to characterize the underlying mechanisms. In particular, we focused on the mean time to fix the first incompatibility allele or haplotype under recurrent mutations, which can be decomposed into the mean waiting time of occurrence of the first mutation destined to be fixed (*T*_*wait*_) and the mean fixation time conditioned on fixation (*T*_*fix*_):

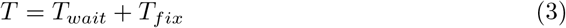

Then we performed multi-loci simulations to assess how the results scale up the genome scale.

### Single-locus incompatibility

#### Underdominant mutations

This model has already been studied by Charlesworth [38] but we summarized it for completeness and provided additional results. We considered a single bi-allelic locus, with the ancestral allele *A*_1_ that can mutate to the derived allele *A*_2_ at rate *µ*. The fitness of genotypes *A*_1_*A*_1_, *A*_1_*A*_2_, and *A*_2_*A*_2_ are 1, 1 *− s*_*u*_, and 1 + *s*, respectively, and the frequency of allele *A*_2_ is noted *x*.

The change in allele frequencies in one generation is given by:

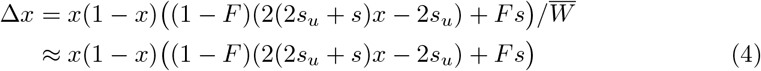

where 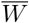 is the mean fitness of the population, approximately equal to 1 when selection is weak. Equation (4) can also be written as:

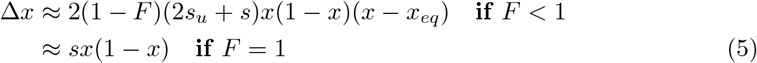

with

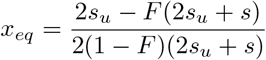

**Table 1.** Glossary of the main notations.

| Symbol | Biological meaning |
| --- | --- |
| $N$ | Population size |
| $N_e$ | Effective population size |
| $\sigma$ | Selfing rate |
| $F$ | Wright's fixation index |
| $\mu$ | Mutation rate |
| $r$ | Recombination rate |
| $L$ | Genome length (used in multi-loci simulation models only) |
| $h$ | Coefficient of dominance (besides genetic incompatibilities) |
| $s$ | Strength of selection (besides genetic incompatibilities) |
| $h_c, h_B$ | Coefficient of dominance of the compensatory and BDMi mutations |
| $s_u, s_c, s_B$ | Strength of selection of the underdominant, compensatory and BDMi mutations |
| $k_c, k_B$ | Coefficient of dominance in double heterozygotes for compensatory and BDMi mutations |

According to diffusion theory, the probability of fixation of a single *A*_2_ mutant is given by:

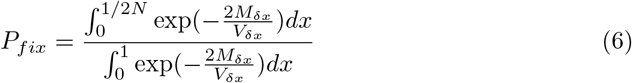

where *M*_*δx*_ = Δ*x* is the expected infinitesimal change in allele frequency, and 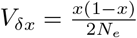 is the expected infinitesimal variance.

#### Two-loci incompatibilities

We also considered models with two bi-allelic loci. *A*_1_ and *A*_2_, *B*_1_ and *B*_2_ denote the ancestral and derived alleles at the first and second locus, respectively, and their respective frequencies are *x* and 1 − *x, y* and 1 − *y*. We assumed the same mutation rate, *µ*, at the two loci. The recombination rate between the two loci is 0 ≤ *r* ≤ 0.5, where 0 corresponds to fully linked loci and 0.5 corresponds to loci located on different chromosomes with Mendelian segregation. The frequency of the four haplotypes *{A*_1_*B*_1_, *A*_1_*B*_2_, *A*_2_*B*_1_, *A*_2_*B*_2_*}* are noted as *{X*_1_, *X*_2_, *X*_3_, *X*_4_*}*, and the frequency of the ten genotypes, the combination of haplotypes *X*_*i*_ and *X*_*j*_, are noted as *G*_*ij*_ with *i ≤ j* (for example *G*_12_ = [*A*_1_*A*_1_; *B*_1_*B*_2_]). Note that we must distinguish *G*_14_ from *G*_23_, which correspond to identical genotypes [*A*_1_*A*_2_; *B*_1_*B*_2_] but may differ in the haplotypes produced through gametogenesis. Changes in genotype frequencies can be obtained as follows [38]:

After meiosis, haplotype frequencies are given by:

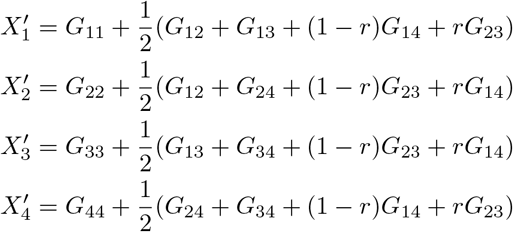

After syngamy, genotypic frequencies are given by:

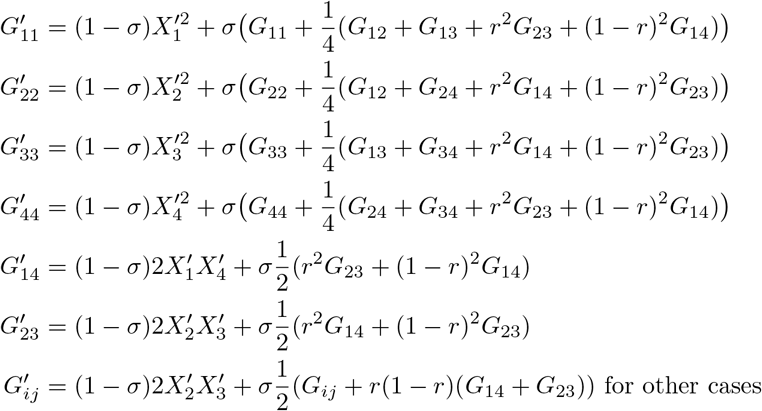

Finally, after selection:

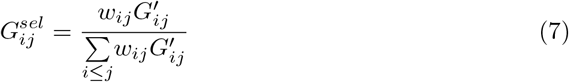

where *w*_*ij*_ is the fitness of genotype *G*_*ij*_, which depends on the type of two-loci incompatibility studied (see Compensatory mutations and BDMi mutations below). Note that in either scenarios, we do not distinguish *cis* and *trans* effects on fitness (*i*.*e*., *w*_14_ = *w*_23_).

#### Compensatory mutations

We extended previous models of compensatory mutations [39, 40] by including the effects of partial selfing. In brief, compensatory mutations at two loci can be viewed as a generalization of the one-locus underdominant model presented above where two haplotypes are equally fit, *A*_1_*B*_1_ and *A*_2_*B*_2_, but the intermediate paths, *A*_1_*B*_2_ and *A*_2_*B*_1_ are deleterious. Thus, alike underdominant mutations, the evolution of pairs of compensatory mutations requires to cross a fitness valley.

In the compensatory mutation models, we set the fitness of genotypes as follows:

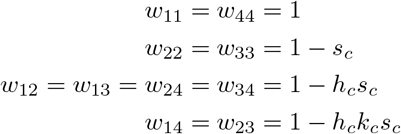

where *s*_*c*_ *≥* 0 and 0 *≤ h*_*c*_ *≤* 1 are the strength and the coefficient of dominance of the deleterious effects of each mutation respectively, and *k*_*c*_ is the coefficient of dominance for the double heterozygous genotype *A*_1_*A*_2_*B*_1_*B*_2_.

#### BDMi mutations

We extended the model of Kimura and King [41] by including the effects of partial selfing. BDMi mutations generate genetic incompatibility only when an individual carries both derived alleles. Thus, individuals carrying either derived alleles do not experience deleterious effects, meaning that the fixation of BDMi mutations does not require to cross a fitness valley.

In the BDMi mutations models, we set the fitness of the genotypes as follows:

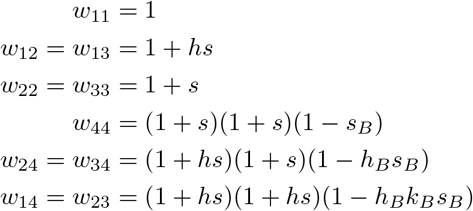

where *s*_*B*_ *≥* 0 and 0 *≤ h*_*B*_ *≤* 1 are the strength and the coefficient of dominance of the genetic incompatibility between the two derived alleles *A*_2_ and *B*_2_ respectively, and *k*_*B*_ is the coefficient of dominance for the double heterozygous genotype *A*_1_*A*_2_*B*_1_*B*_2_. *s ≥* 0 and 0 ≤ *h ≤*1 are the strength of selection and the coefficient of dominance of the local adaptation. For the standard BDMi model *s* is set to 0, and *s >* 0 only when we assume that BDMi mutations are also driven by local adaptation.

### Simulations

We also developed, first, single-locus (underdominance mutations) and two-loci (compensatory and BDMi mutations) simulation models to check for analytical and numerical predictions and, second, multi-loci simulation models (underdominant, compensatory, and BDMi mutations) to test for the possible effect of interactions between segregating mutations.

#### Single-locus and two-loci simulations

We developed C++ programs to simulate populations of *N* hermaphroditic individuals producing gametes with mutations from ancestral to derived alleles at a rate *µ* and, for the two-loci simulations, recombinations between the two loci at a rate 0 ≤ *r* ≤ 0.5. Syngamy may occur through self-fertilisation, at a rate 0 ≤ *σ* ≤ 1. Then, selection on offspring occurs differently in the underdominant, compensatory, and BDMi mutations simulations (as described above). Finally, genetic drift was included by sampling offspring from the genotype frequencies using a multinomial distribution using the *gsl ran multinomial* function from the GNU Scientific Library [42]. For each iteration, we measured the number of generations required to fixate the derived allele (underdominant mutation), a pair of derived alleles (compensatory mutations), and either derived allele (BDMi mutations).

Importantly, selfing is known to increase the strength of background selection, which further reduces effective population size [26, 27, 43]. The strength of this effect critically depends on genomic recombination rate. When the genomic recombination rate is high, only highly selfing populations suffer from background selection. In contrast, when the genomic recombination rate is low, the effect of background selection rise linearly with selfing [26, 27]. To account for this effect, we used a Dirichlet-multinomial distribution in the genetic drift function (instead of a multinomial distribution) that allowed us to tailor the effective size of the population to its selfing rate. For this we used analytical approximations in Roze [27] to simulate two background selection scenarios corresponding to two levels of genomic recombination. The first scenario corresponds to a low rate of genomic recombination and leads to a linear decrease in *N*_*e*_ with selfing rate (hereafter ‘linear BG effects’). The second scenario corresponds to a high rate of genomic recombination and leads to an accelerating decrease in *N*_*e*_ with selfing rate (hereafter ‘curved BG effects’). Specifically, using the BS1 function from the supplemental material of Roze [27] (File S2), we modelled the background selection effects assuming that deleterious alleles with selection and dominance coefficients of *s* = 0.05 and *h* = 0.2 occurred with a genomic mutation rate of *U* = 0.1 in a genome map length of either *R* = 0.5 (’linear BG effects’) or *R* = 40 (’curved BG effects’). This informed us by how much *N*_*e*_ was reduced due to background selection for different selfing rates. Then, we used these background selection effects on *N*_*e*_ to compute the parameter vector *α* of the Dirichlet-multinomial distribution given that:

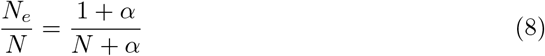

#### Multi-loci simulations

We used individual based forward simulations in which the individuals have diploid genomes of length *L* elements representing loci on which mutations can occur at rate *µ* and between which recombination can occur at rate *r*. As above, we modelled a single population of *N* individuals reproducing through selfing at rate 0 ≤ *σ* ≤ 1 and on which selection depends on the type of genetic incompatibility considered (see above).

The single-locus and two-loci simulation models were performed using C++ scripts, and the multi-loci simulations models were performed on the software SLiM [44], both using GNU parallel [45] (see Code availability).

## Results

### Underdominant mutations

When underdominant, a mutation arising in a population is first counter-selected and it is well known that either genetic drift or selfing facilitates crossing the fitness valley [38]. For completeness we first summarize previous results (Fig. S1 Appendix S1). When selfing rate increases, heterozygous individuals become rarer in the population and so does the selection against underdominant mutations, increasing the probability of fixation and decreasing the time to fixation. When *s >* 0, a new underdominant mutation arising in a population may be directly positively selected if selfing rate is high enough, that is if:

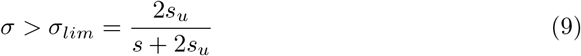

Including the effects of background selection slightly shortens the time to fixation of symmetrical underdominant mutations (*s* = 0), but substantially increases the time to fixation of asymmetrical mutations (*>* 0) in highly selfing populations (Fig. S1 E). This is because background selection reduces the effective population size, *N*_*e*_, and thereby the efficacy of selection. Such a low efficacy of selection virtually reduces both the fitness valley in the heterozygotes, *A*_1_*A*_2_ and the fitness peak in the homozygotes, *A*_2_*A*_2_. But, because background selection mostly impact highly selfing populations – in which selection on heterozygotes is irrelevant – the effects of background selection mostly manifests as a lower probability of fixation in highly selfing populations because the effect of positive selection on the homozygotes is reduced.

Previous studies have only considered a single mutation with constant effect [38]. Interesting properties can be obtained if we assume a distribution of deleterious effect. Assuming that 4*Ns*_*u*_ follows a Gamma distribution with mean *γ* and shape *β*, the proportion of underdominant mutations that can fixate in partially selfing populations compared to outcrossing populations is well approximated by:

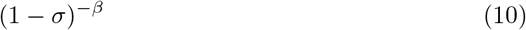

which is independent of *γ*. Thus, because the time to fixation is mostly determined by the waiting time that a mutation that will get fixed arises in a population, the relative time to fixation in partially selfing compared to outcrossing populations is simply given by:

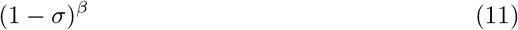

which our multi-loci simulation results confirm as long as *γ* is not too small (Fig. 1). It shows that when *β <* 1 there is a non-linear accelerating effect of selfing on accumulation of underdominant mutations and the lower *β* the stronger the effect.

**Fig 1.**
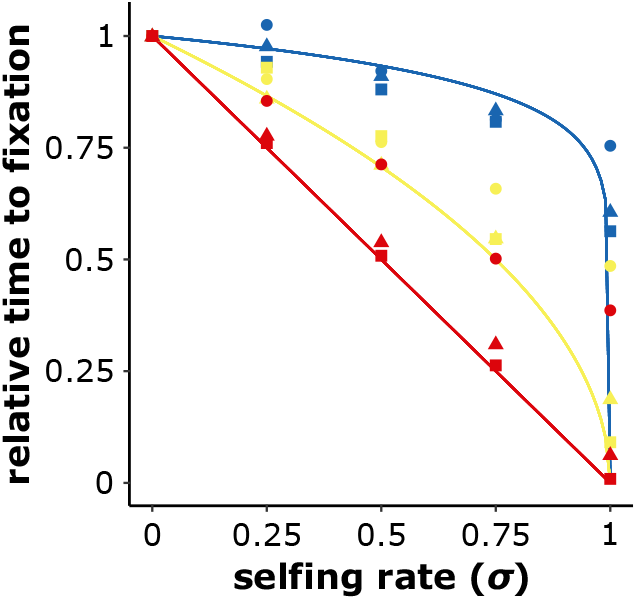
Underdominant mutations accumulate more rapidly in selfing populations. Analytical predictions (lines) and outcomes of multi-loci simulations (symbols) of the averaged number of generations required to fixate a symmetrical underdominant mutation compared to an outcrossing population (*σ* = 0). 4*Ns*_*u*_ follows a Gamma distribution with a shape parameter *β* of 0.1 (green), 0.5 (blue), or 1 (red), and mean parameter *γ* of 10 (circles), 100 (triangles), or 1,000 (squares). *L* = 100, *N* = 1, 000, *µ* = 10^*−*6^, *r* = 0.01. The analytical predictions are (1 *− σ*)^*β*^. 1, 000 *iterations*.

### Compensatory mutations

Compensatory mutations can also contribute to RI, and their fixation also requires crossing a fitness valley. How selfing affects the fixation of such mutations depends first on the strength of the deleterious effect relative to drift (*N*_*e*_*s*_*c*_ of the order of 1 or lower) (Fig. 2). When it is low, fixation occurs in two steps. First, a primary mutation goes to fixation as a weakly deleterious mutation and, second, the compensatory mutation restoring fitness goes to fixation as a weakly beneficial mutation. As the total time mainly depends on the waiting time for mutations that will be fixed arise, it can be approximated by:

**Fig 2.**
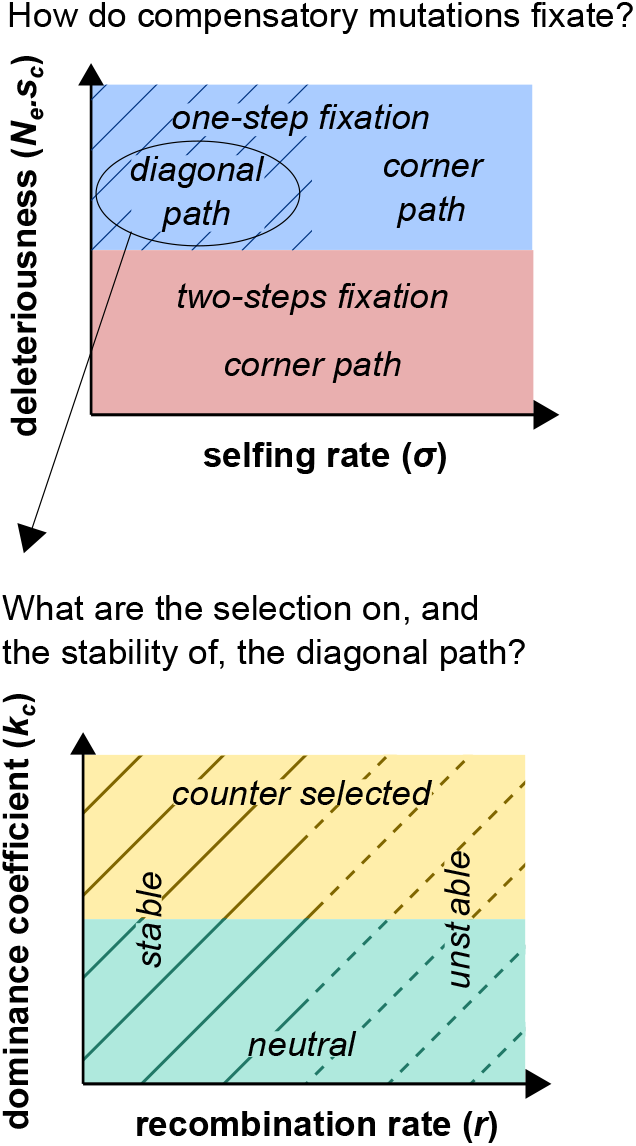
Fixation regime of compensatory mutations as a function of the selfing rate (*σ*), the strength of deleterious effect (deleteriousness), the recombination rate between the two loci (*r*), and the coefficient of dominance in double heterozygotes (*k*_*c*_). Populations with high selfing rates always take the corner path on the fitness landscape (*i*.*e*., the deepest point of the fitness valley represented by genotypes that are homozygous for the ancestral allele at one locus, and homozygous for the derived allele at the other locus), while populations with low selfing rates can take the diagonal path (*i*.*e*., double heterozygous genotype). The diagonal path may however be counter selected (when *k*_*c*_ is relatively high) and unstable (when the recombination rate is relatively high). Note that, for the sake of simplicity, all effects are here presented as qualitative categories while they are in fact gradual.

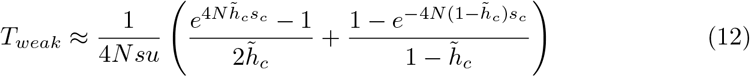

where 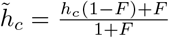, which corresponds to the effective dominance level (*h*_*c*_(1 − *F*)+*F*) scaled by the increase in drift due to selfing (1 + *F*). The factor 2 in the first step corresponds to the two potential initial paths.

Selfing may affect both steps but, depending on the coefficient of dominance (*h*), fixation is either faster or slower in selfing populations than in outcrossing population because highly selfing populations are less affected by the selection at the heterozygous state (Fig. S2). For instance, recessive deleterious mutations (*e*.*g*., *h*_*c*_ = 0), do not experience fitness costs in heterozygotes, which facilitates the fixation of both the primary deleterious and the secondary beneficial mutations in outcrossing populations [46]. In contrast, dominant deleterious mutations (*e*.*g*., *h*_*c*_ = 1), do experience fitness costs in heterozygotes, which both hamper the fixation of the primary deleterious mutation and hide the beneficial effects of the second one in outcrossing populations. Overall, selfing speeds up the fixation of compensatory mutations if the mutation is dominant (*h*_*c*_ *>* 1*/*2), and slows it down if the mutation is recessive (*h*_*c*_ *<* 1*/*2), and has no effects when mutation is codominant (*h*_*c*_ = 1*/*2 hence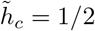, independent of *F*) (Fig. S2).

In contrast, when the deleterious effect is too high (*N*_*e*_*s*_*c*_ *>>* 1) to allow the fixation of a singly mutation, then the two compensatory mutations must segregate together in the population and fixate together. In this case, the key parameters are the recombination rate between loci (*r*), and the coefficient of dominance in the double heterozygote (*k*_*c*_) (Fig. 2). When *r* = 0 and *k*_*c*_ = 0, we found that the fixation time of a pair of compensatory mutations (*T*_0,0_) may be approximated by (see Appendix S1):

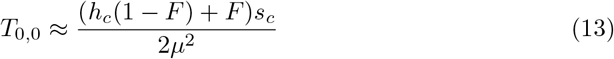

which shows that selfing always increases the fixation time (Fig. 3B and Fig. S3). This is because evolution can follow the diagonal path of the fitness landscape (*i*.*e*., through the double heterozygous genotype *A*_1_*A*_2_*B*_1_*B*_2_), and thus avoid the corner path made of genotypes with low fitness value (*i*.*e*., *A*_1_*A*_1_*B*_2_*B*_2_ or *A*_2_*A*_2_*B*_1_*B*_1_), more easily in outcrossing than in selfing populations (Fig. 2, Fig. 3, Fig. S4). In comparison to the corner path, the diagonal path represents a lower fitness valley, which flattens out as *k* approaches 0. Thus, when *r* and *k*_*c*_ are both equal to 0, outcrossing populations fixate compensatory mutations by forming the double mutated haplotype (*A*_2_*B*_2_), which is stable because it is not broken down by recombination and can spread in the population because it is less counter selected.

**Fig 3.**
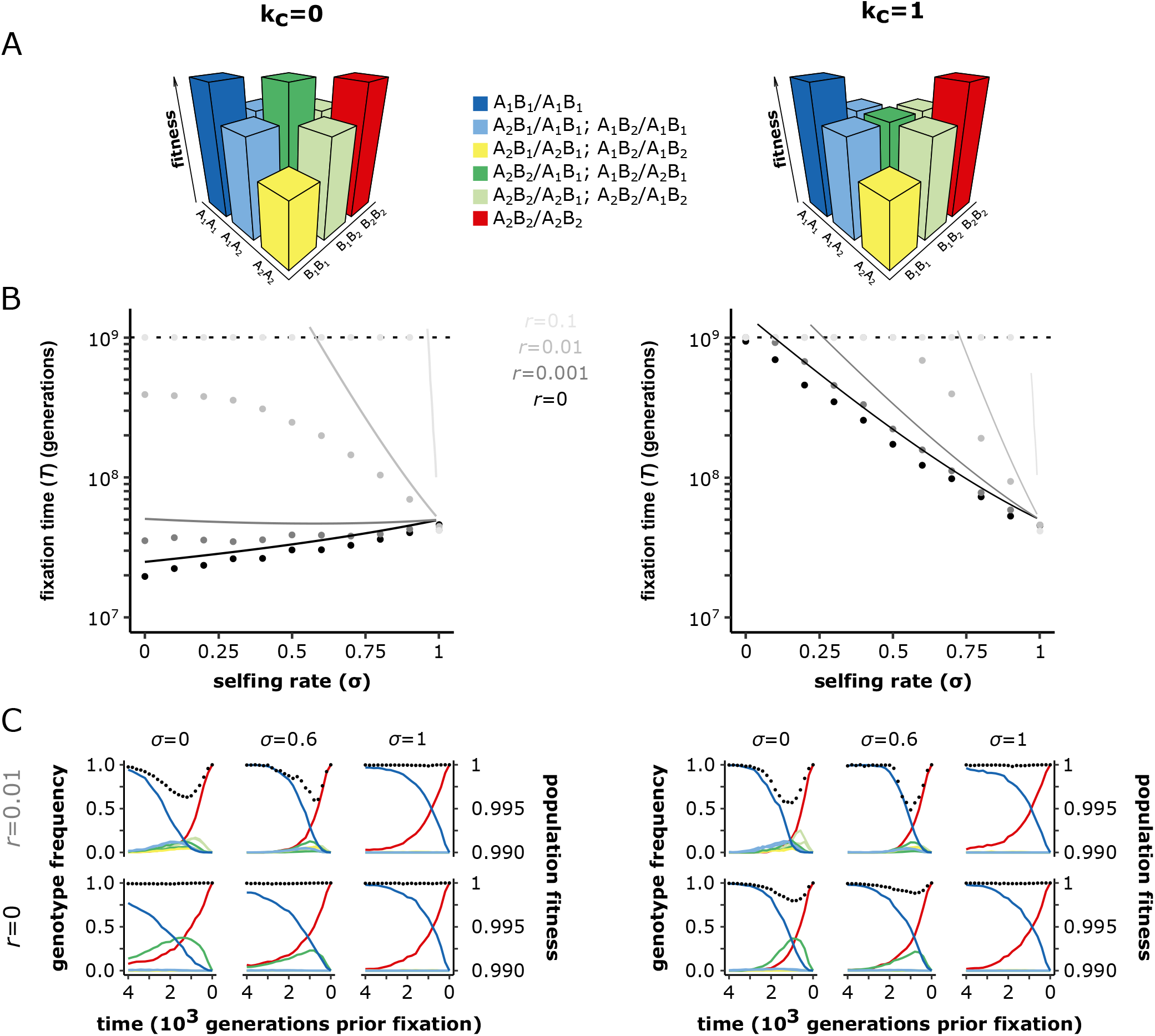
The effect of selfing on the fixation of compensatory mutations depends on the coefficient of dominance in the double heterozygote (*k*_*c*_), and the recombination rate(*r*). (A) Schematic representations of the fitness landscapes for compensatory mutations with *k*_*c*_ = 0 (left) or *k*_*c*_ = 1 (right). *A*_1_ and *B*_1_ are the ancestral alleles. *A*_2_ and *B*_2_ are the derived alleles. Time to fixation of a pair compensatory mutations with *k*_*c*_ = 0 (left) or *k*_*c*_ = 1 (right). The lines correspond to the analytical approximations (eq. 14) for *r* = 0 (black), *r* = 0.001 (dark grey), *r* = 0.01 (grey) and *r* = 0.1 (light grey). Dots show the corrected mean times to fixation of a pair of compensatory mutations from our two-loci simulations. We corrected the raw means because the threshold made our data right censored (*i*.*e*., missing estimates above 10^9^ generations), which we accounted for by first estimating the full distribution by fitting gamma distributions on our simulation outputs using the *fitdistriplus* R package [47], from which we estimated the corrected mean times to fixation. The dashed horizontal lines indicate the generation threshold after which simulations stop. *N* = 1, 000, *µ* = 10^*−*5^, *h*_*c*_ = 0.5, *s*_*c*_ = 0.01. 1, 000 *iterations*. (C) Population fitness (black dots – right y axis) and the frequencies of the 10 possible genotypes on the two loci fitness landscapes (solid lines – left y axis) over the last 4200 generations preceding the fixation of the pair of compensatory mutations. The line colours match the genotype colours on the fitness landscapes. *N* = 1, 000, *µ* = 10^*−*5^, *h*_*c*_ = 0.5, *s*_*c*_ = 0.01. 100 *iterations*. See Fig. S3 and Fig. S4 for the visualisation of additional parameter combinations.

When *k*_*c*_ *>* 0 and/or *r >* 0, the dynamics is very different. Either selection against double heterozygotes or recombination that breaks down double-mutated haplotypes considerably reduces the probability of crossing the fitness valley. When *r* is small, we can show that:

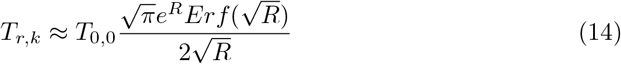

where *Erf* is the error function and *R* = *N* (1 − *σ*)(*r* + *hks*). The right-hand function increases very rapidly with *R*. These approximations and our simulations show that both recombination and selection against heterozygotes strongly impedes the fixation of compensatory mutations, both effects being reduced by selfing (Fig. 3B and Fig. S3). When *k*_*c*_ *>* 0, the diagonal path represents a fitness valley that selects against the double mutated haplotype (*A*_2_*B*_2_). And, when *r >* 0, the double mutated haplotype (*A*_2_*B*_2_) may break, forming genotypes that are off the diagonal path and eliminating the derived alleles (*A*_2_ and *B*_2_) (Fig. 2). In contrast, the fixation time in highly selfing populations is less influenced by *r* and *k*_*c*_ because their high level of homozygosity (i) makes selfing populations not following the diagonal path of the fitness landscape, and (ii) makes the double mutated haplotype (*A*_2_*B*_2_) more stable over time due to the inefficient recombination in selfers [23].

We further found that, regardless of the coefficient of dominance in the double heterozygote (*k*_*c*_) and the recombination rate (*r*), background selection shortens fixation time in highly selfing populations (Fig. S5). By increasing drift, background selection reduces the efficacy of selection against the primary deleterious mutation, which may thus segregate at higher frequency and be more likely to be associated with the secondary compensatory mutation. Moreover, because of background selection, *N*_*e*_*s* can be small enough under selfing for the fixation to occur in two steps (which is rapid see (13)) whereas, for the same population size, *N*_*e*_*s* can be too high under outcrossing so that fixation can occur in a single step (which is much longer (14)).

The outcome of our two-loci simulations concur with the multi-loci simulations. Specifically, our multi-loci simulations indicate that the fixation of two compensatory mutations depends on the coefficient of dominance in the double heterozygote (*k*_*c*_), and the recombination rate (*r*). Selfing speeds up the fixation of compensatory mutations, except when the fitness of the double heterozygote is neutral (*k*_*c*_=0) and the recombination rate is low enough (*r <* 0.0001) for the haplotype carrying both derived alleles to be stable (Fig. S6).

### BDMi mutations

Contrary to our models of underdominant and compensatory mutations, the BDMi model does not require crossing a fitness valley for RI to accumulate. Therefore, there is no obvious reason for selfing to accelerate RI. We first consider mutations that behave neutrally in isolation (*s* = 0). If mutation rate is low compared to drift (4*N*_*e*_*µ <* 1), the two incompatible alleles (*A*_2_ and *B*_2_) rarely segregate at the same time in the population and one can consider that they fix independently. According to basic results of the neutral theory, Equation 3 reduces to:

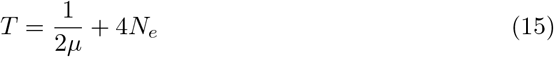

The 2*µ* term comes from the fact that mutation can occur at the two loci. In this case, selfing only shortens the time a mutation needs to spread through the population and get fixed (4*N*_*e*_), but does not affect the waiting time (1*/*2*µ*). Because the former may be negligible compared to the latter (as implicitly assumed in [37]), selfing barely impacts the fixation time as confirmed by simulations (see below).

However, when the mutation rate is high (4*N*_*e*_*µ >* 1), the mean time a mutation needs to get fixed is shorter than 4*N*_*e*_ because mutations of the same type may arise on different individuals of the populations, so that multiple mutations can get fixed simultaneously. Taking this effect into account, Kimura [48] showed that, at a single locus, the mean time to fixation under continuous mutation pressure was:

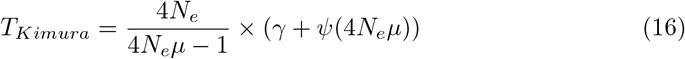

where *γ* is Euler’s constant and *ψ* is the digamma function. Note that (16) converge to (15) for small *N*_*e*_*µ*. Because our BDMi models have two mutation types (for the A and B loci), we cannot use equation (16) as such because mutations of different types cannot get fixed simultaneously in the populations. As a heuristic argument, we can decompose *T*_*Kimura*_ = *T*_*wait*_ + *T*_*fix*_ as in equation (3), where *T*_*wait*_ = 1*/µ*. Thus, by using *T*_*fix*_ = *T*_*Kimura*_ *−* 1*/µ* instead of 4*N*_*e*_ in (15), a more accurate expression under high mutation rate is:

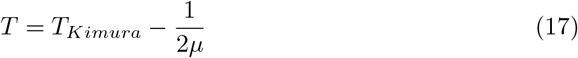

However, mutations can segregate at the two loci at the same time and be jointly selected against, which is not taken into account in (17). Equation 17 thus serves as a neutral reference to assess whether incompatibilities between mutations segregating within populations affect the fixation time of BDMi mutations, and whether it does so independently of the population selfing rate. Simulations showed that several BDMi mutations could often segregate together in the population (Fig. S8), and cause incompatibilities within populations (Fig. 5A, Fig. S7). Accordingly, simulations showed that high mutation rate and/or high effective size slowed down the fixation of BDMi mutations (Fig. 4). Counter selection of such segregating incompatibilities hamper the fixation of either derived alleles, and stronger (high *s*_*B*_; Fig. 6A) or more dominant incompatibilities (*h*_*B*_ *>* 0.5; Fig. 6B) increase fixation time. Finally, recombination helps forming incompatible *A*_2_*B*_2_ haplotypes and also reduces the accumulation of BDMi mutations (Fig. 4). Thus, selfing has opposing effects on these dynamics. On the one hand, it increases selection by exposing the incompatible haplotype in homozygouste state (especially when *h*_*B*_ is low). On the other hand it increases drift and reduces genetic shuffling, which reduce the occurrence of the incompatible haplotype. Overall the second effects dominate and selfing globally reduces the time to fixation of BDMi mutations.

**Fig 4.**
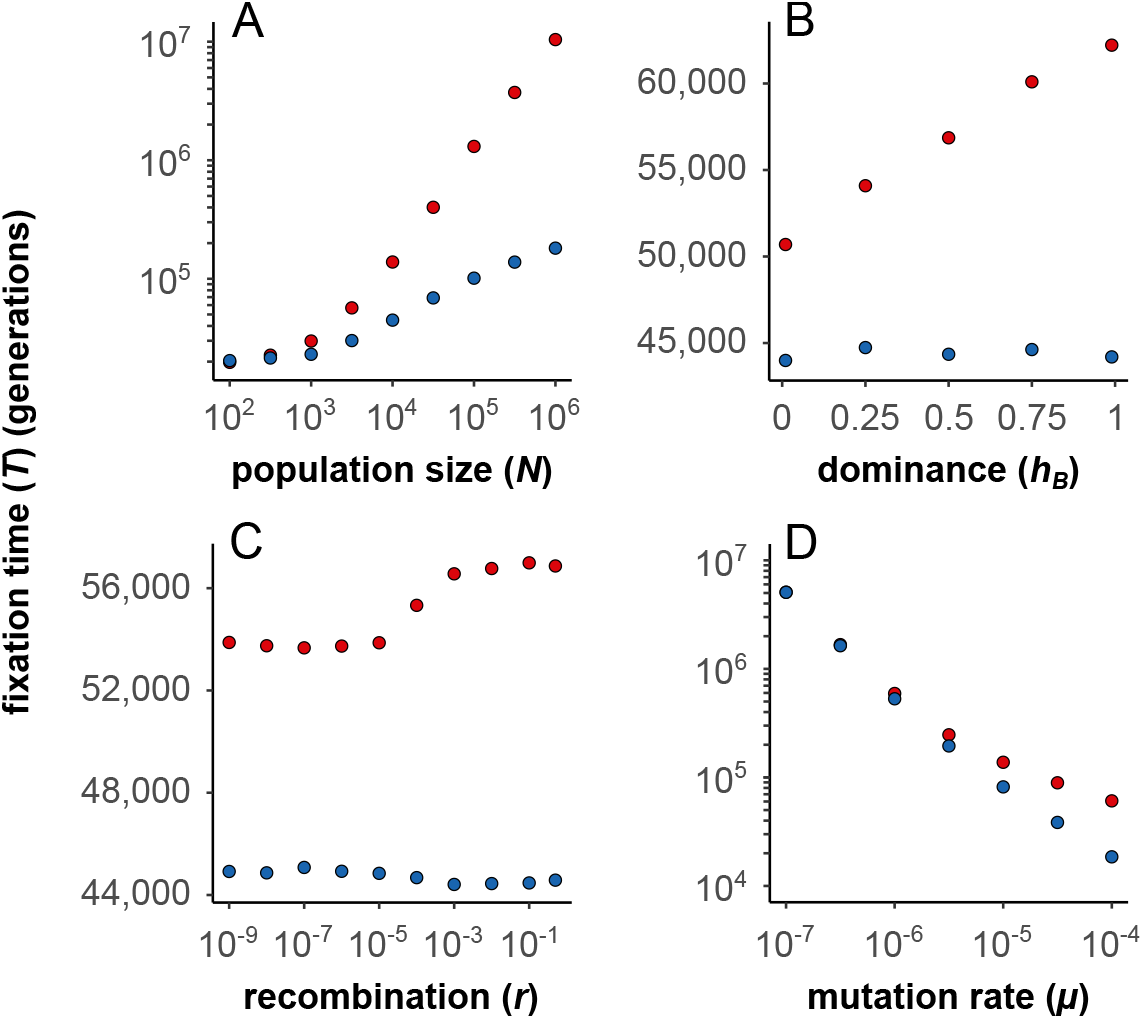
Fixation time of BDMi mutations (red) compared to neutral mutations (blue) in outcrossing populations (*σ* = 0) (two-loci model). (A) Higher population sizes (N) decelerate the fixation of BDMi mutations (*s*_*B*_ = 2.5.10^*−*2^, *h*_*B*_ = 0.5; red) compared to neutral mutations (*s*_*B*_ = 0; blue). *r* = 0.5, *µ* = 2.5.10^*−*5^. 1, 000 *iterations*. (B) Higher coefficients of dominance (*h*_*B*_) decelerate the fixation of BDMi mutations (*s*_*B*_ = 2.5.10^*−*4^; red) compared to neutral mutations (*s*_*B*_ = 0; blue). *N* = 10, 000,*r* = 0.5, *µ* = 2.5.10^*−*5^. 10, 000 *iterations*. (C) Higher rates of recombination (*r*) decelerate the fixation of BDMi mutations (*s*_*B*_ = 2.5.10^*−*4^, *h*_*B*_ = 0.5; red) compared to neutral mutations (*s*_*B*_ = 0; blue). *N* = 10, 000, *µ* = 2.5.10^*−*5^. 100, 000 *iterations*. (D) Higher rates of mutation (*µ*) accelerate the fixation of BDMi mutations (*s*_*B*_ = 2.5.10^*−*3^, *h*_*B*_ = 0.5; red) compared to neutral mutations (*s*_*B*_ = 0; blue). *N* = 10, 000, *r* = 0.5. 10, 000 *iterations*.

**Fig 5.**
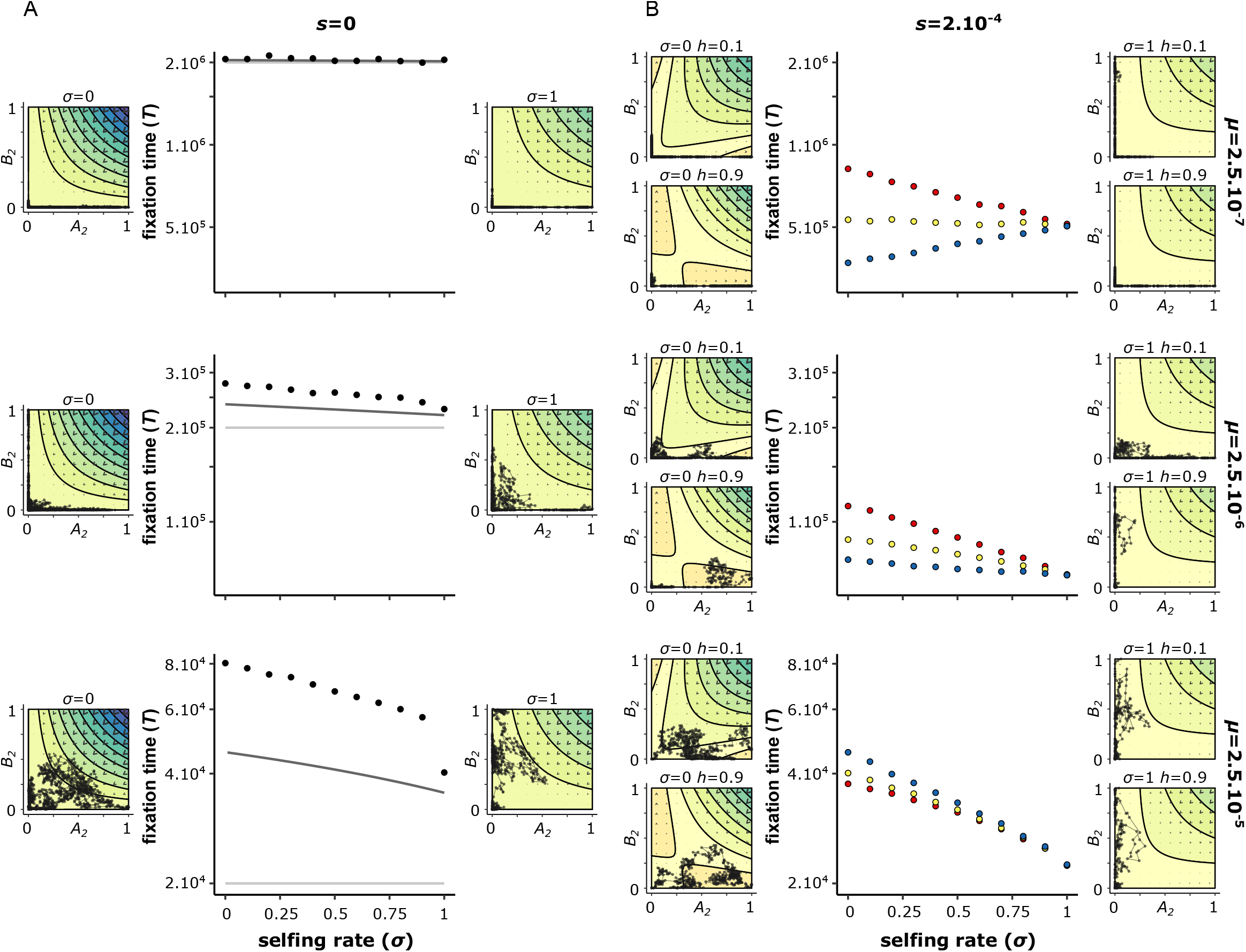
Selfing rate accelerates the fixation of BDMi mutations for high mutation rates. The fixation time estimated from the two loci models with mutation rates (*µ*) for *A*_2_ and *B*_2_ of either 2.5.10^*−*7^ (top), 2.5.10^*−*6^ (middle), or 2.5.10^*−*5^ (bottom), and the strength of selection on the derived alleles (*s*) is either 0 (left), or 2.10,^*−*4^ (right). When there is selection, the coefficient of dominance (*h*) is either recessive (*h* = 0.1, red), co-dominant (*h* = 0.5, yellow), or dominant (*h* = 0.9, blue). The solid lines in the neutral scenario correspond to analytical approximations: 1*/*2*µ* (light grey), and equation (17) (dark grey) (see BDMi Results section for details on the approximations). Each phase portrait shows, for a single simulation, the change in allele frequencies of *A*_2_ and *B*_2_ plotted from the beginning, and then every 100 generations until the fixation of a derived allele. The isoclines represent the expected benefits (warm colour) and costs (cold colours) on population fitness (multiplied by *N*_*e*_, and with a increment of 1). The direction of the arrow indicate the expected allele change (which is the balance between mutation rates and selection), and their size indicates the strength of the change. *N* = 10, 000, *h*_*B*_ = *k*_*B*_ = 0.5, *s*_*B*_ = 10^*−*3^, *r* = 0.5. 10, 000 *iterations*.

**Fig 6.**
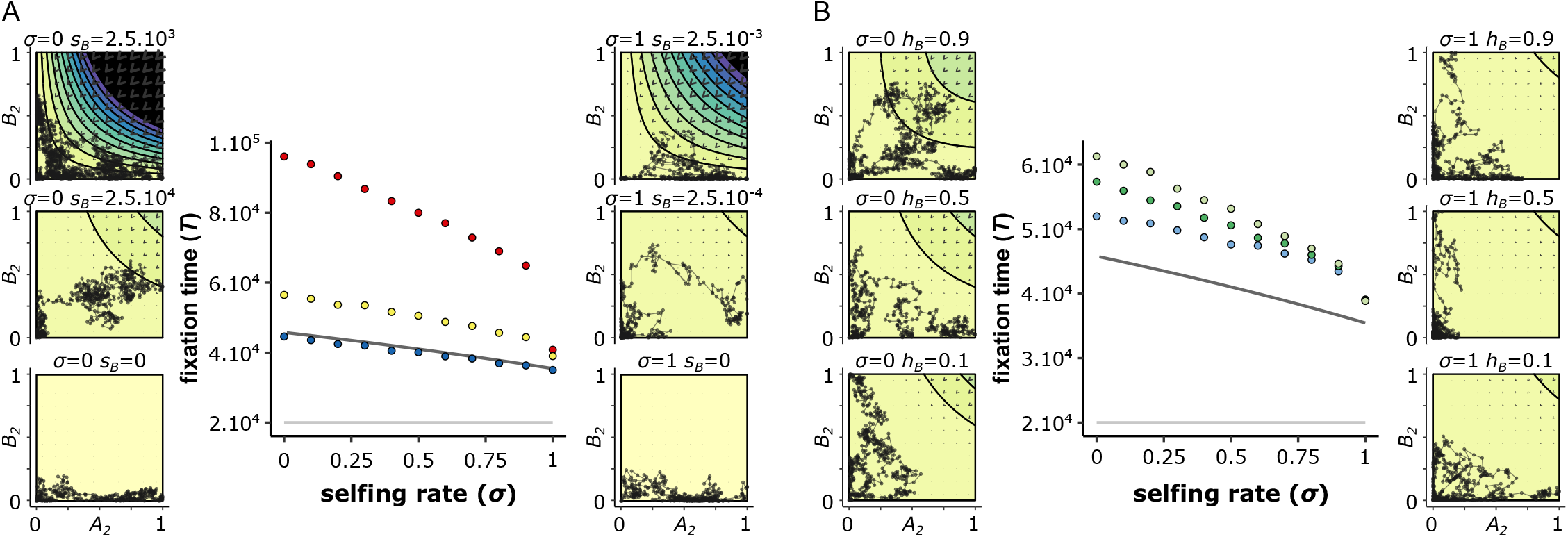
Within-population interferences between BDMi mutations slow down their fixation, and occur less in highly selfing populations (two loci model). The A panel shows the fixation time of co-dominant BDMi mutations (*h*_*B*_ = 0.5), and with a strength of selection (*s*_*B*_) of either 0 (blue), 2.5.10^*−*4^ (yellow), or 2.5.10^*−*3^ (red). The B panel shows the fixation time of BDMi mutations with a strength of selection (*s*_*B*_) of 2.5.10^*−*4^ that are either recessive (*h*_*B*_ = 0.1; light blue), co-dominant (*h*_*B*_ = 0.5; green), or dominant (*h*_*B*_ = 0.9; light green). The solid lines in the neutral scenario correspond to analytical approximations: 1*/*2*µ* (light grey), and equation (17) (dark grey) (see BDMi Results section for details on the approximations). Each phase portrait shows, for a single simulation, the change in allele frequencies of *A*_2_ and *B*_2_ plotted from the beginning, and then every 100 generations until the fixation of a derived allele. The isoclines represent the expected benefits (warm colour) and costs (cold colours) on population fitness (multiplied by *N*_*e*_, and with a increment of 1). The direction of the arrow indicate the expected allele change (which is the balance between mutation rates and selection), and their size indicates the strength of the change. *N* = 10, 000, *k*_*B*_ = *h*_*B*_, *s* = 0, *r* = 0.5. 10, 000 *iterations*.

When there is direct selection on mutations (*s >* 0), the effect of selfing on the accumulation of BDMi mutations depends on the interaction between the mutation rate (*µ*) and the coefficient of dominance (*h*) (Fig. 5B, Fig. S7). When the mutation rate is low, BDMi mutations get fixed like beneficial mutations, whose probability of fixation depends on selfing rate and the coefficient of dominance (*h*) [46]. When the mutation is recessive (*i*.*e*., *h <* 0.5), the probability of fixation increases with selfing. When the mutation is dominant (*i*.*e*., *h <* 0.5), the probability of fixation decreases with selfing. And, when the mutation is codominant (*i*.*e*., *h* = 0.5), selfing does not affect the probability of fixation [46]. Therefore, when mutation rate is low, selfing speeds up fixation of recessive BDMi mutations, and slows down fixation of dominant BDMi mutations and can be approximated by equation (13) in [46] (without standing variation, *P*_*sv*_ = 0, their notations) (Fig. 5B, Fig. S7). In contrast, when mutation rate increases, selfing speeds up fixation of BDMi mutations regardless of the coefficient of dominance (*h*). This is because it is more likely that multiple BDMi mutations segregate in the population (Fig. S8) and cause genetic incompatibilities within populations, hampering more the fixation of BDMi mutations in outcrossing population than in selfing populations, as described above.

The fixation time of BDMi mutations is also affected by background selection, which has two main effects: higher drift makes selection less efficient when selfing increases and the occurrence of segregating mutations at both loci less likely. When there is no direct selection (*s* = 0), BDMi are less counter selected within populations and accumulate faster under selfing than under outcrossing. So, background selection reinforces the effect of selfing. When there is direct selection (*s >* 0) selfing reduces both selection on the beneficial allele and against the incompatible haplotype. When mutation rate is low and selection against the incompatible haplotype limited even in outcrossing populations, reducing selection on the beneficial allele predominate and selfing slows down the accumulation of BDMi mutations (Fig. 7, Fig. S9). However, for higher mutation rates, reduced selection against the incompatible haplotype predominates and selfing speeds up fixation. Overall, under a wide range of conditions, even with direct selection, selfing facilitates RI.

**Fig 7.**
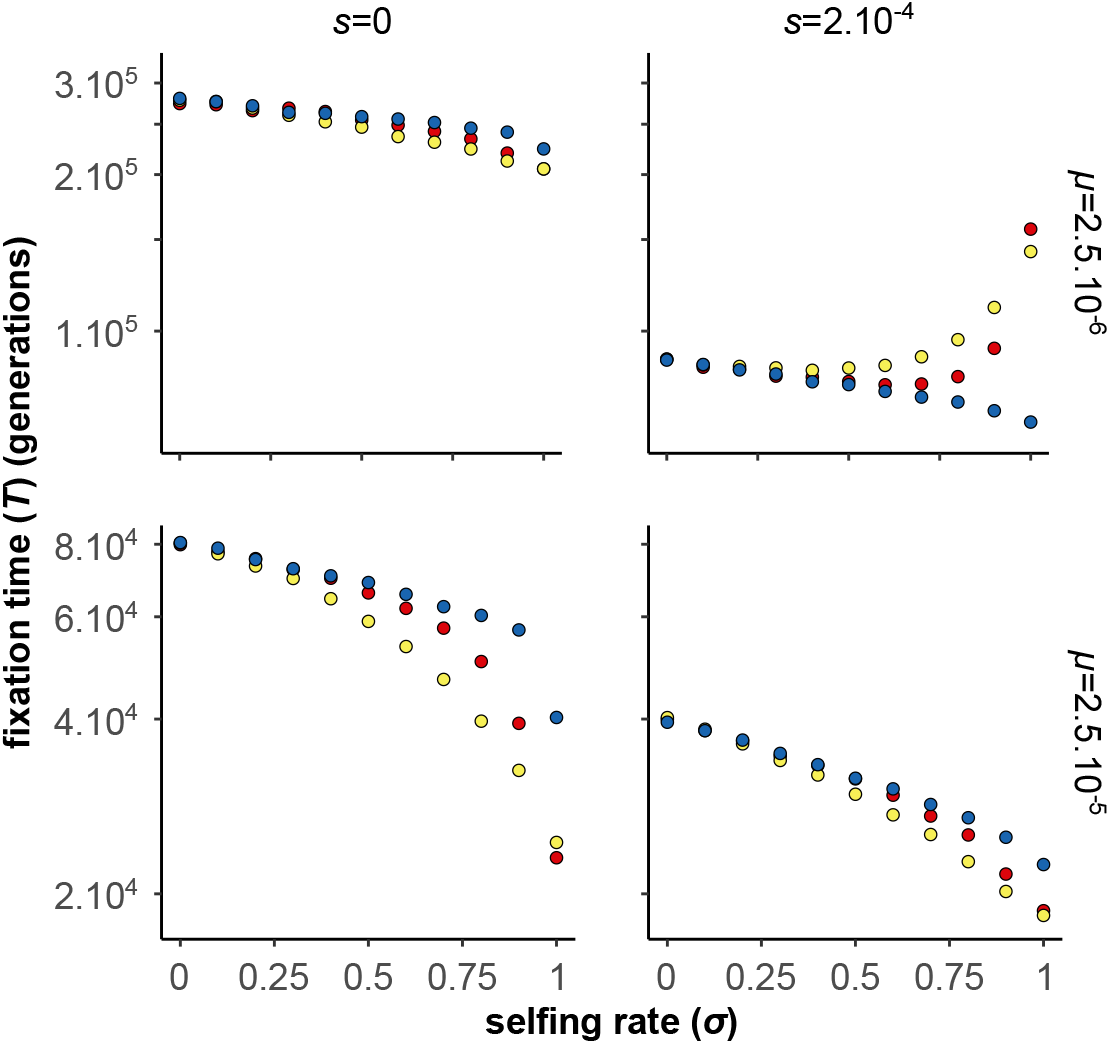
Effects of background selection on the accumulation of BDMi mutations (two-loci model). The graph displays the fixation time, either without (left) or with (right) selection on the derived alleles, and under different scenarios of background selection, ‘linear BG effects’ (yellow) or ‘curved BG effects’ (red), which we compared to a scenario without background selection (blue) (see methods for details on the implementation of background selection effects in our simulations). *N* = 10, 000, *h* = 0.5, *s* = 2.5.10^*−*4^, *h*_*B*_ = *k*_*B*_ = 0.5, *s*_*B*_ = 10^*−*3^, *r* = 0.5. 10, 000 *iterations*.

In the previous analyses we considered that the direct selection on the *A*_2_ or *B*_2_ alleles is independent of selfing (*s* is a constant), as for adaptation to a new environment. Selfing only modulated the efficacy of selection through its effect on *N*_*e*_ and homozygosity. However, selfing can also directly affects selection, in particular in case of genetic or sexual conflicts [49]. If such conflicts play an important role in RI, it has been proposed that selfing may slow down speciation [9]. We did not explore an explicit model involving conflicts, but to mimic such a situation we considered the simple case where selfing directly reduced the selection coefficient: *s* = *s*_0_(1 − *σ*). This is a strong effect as selection vanishes under full selfing. We consider the case of high mutation rate with background selection that corresponds to the best conditions under which selfing promotes speciation. If “conflict” selection is weak (*s*_0_ here), selfing still facilitates the accumulation of BDMi. However, for stronger selection, despite negative interaction among segregating BDMi, fixation of BDMi is faster under outcrossing than selfing (Fig. 8).

**Fig 8.**
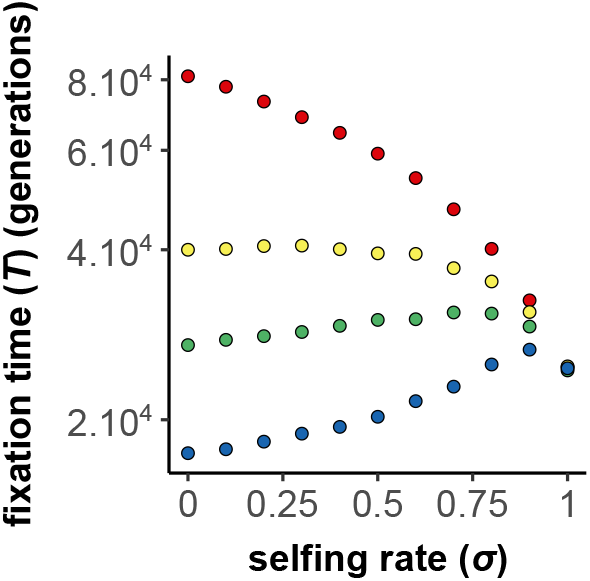
Effects of selfing on the fixation time of BDMi mutations when the strength of selection (*s*) varies with selfing rate (*σ*) (two-loci model). The strength of selection (*s*) decreases with selfing rate as follows: *s* = *s*_0_(1 − *σ*), with *s*_0_ = 0 (red), *s*_0_ = 2.10^*−*4^ (yellow), *s*_0_ = 4.10^*−*4^ (green), and *s*_0_ = 8.10^*−*4^ (blue). *N* = 10, 000, *µ* = 2.5.10^*−*5^, *h* = 0.5, *h*_*B*_ = *k*_*B*_ = 0.5, *s*_*B*_ = 10^*−*3^, *r* = 0.5. 10, 000 *iterations*.

## Discussion

The role of mating systems in speciation is an old question, in particular among plant evolutionary biologists [1, 50, 51]. Depending on the underlying mechanisms, selfing has been proposed to either promote or hamper speciation [8, 9]. Surprisingly, despite this long-standing interest, specific models on the role of selfing on RI are scarce and mainly concerns the build-up of RI caused by underdominant mutations [38]. We filled this gap by expanding previous theoretical work on underdominant mutations and by considering RI caused by epistatic mutations, such as compensatory mutations and Bateson-Dobzhansky-Muller incompatibility mutations. Overall, we showed that selfing promotes allopatric speciation for a wide range of parameters. In addition, our results predicts that mating systems should affect the genomic architecture of reproductive isolation.

### Selfing helps crossing fitness valleys

The Bateson-Dobzhansky-Muller model was initially proposed as a possible solution of the puzzling question of the evolution of hybrid incompatibilities as it does not require crossing fitness valleys [52]. Alternatively, some mechanisms can help crossing fitness valleys, and it is well known that selfing facilitates the fixation of underdominant mutations [38]. We extended this model to a two-loci fitness landscape for which selfing also helps crossing the valley under most conditions. Selfing has two main effects: first it increases drift (in particular if background selection is strong); second, because of reduced recombination and heterozygozity, it limits the breakdown of the new fittest genotype, once it has been created. At the genome scale, assuming several loci where underdominant fitness landscape may occur, we also showed that the effect of selfing is stronger when there is a highly skewed distribution of the depth of the valleys to be crossed (11). As similar conclusion is also likely for the compensatory mutation model although we did not obtain an equivalent analytical result.

### The role of interferences among mutations segregating within populations

In the simplest form of the Bateson-Dobzhansky-Muller model of speciation, genetic incompatibilities occur between derived alleles that are supposed to arise and become fixed independently in different populations [7, 53], and the phase during which BDMi mutations emerge, spread through the population and eventually get fixed is often dismissed (e.g. [36]). This inevitable phase has recently been argued to have important implications in speciation genetics [54]. Considering that BDMi alleles may segregate in natural populations at polymorphic frequencies allows for instance to better explain why hybrid incompatibility may be variable between different pairs of individuals originating from the same two populations (reviewed in [54]), and (ii) why genetic incompatibilities is widespread within species, as found in *Drosophila melanogaster* [55], *Caenorhabditis elegans* [56], *Arabidopsis thaliana* [57], or in the *genus Draba* [18].

To mimic genetic incompatibilities arising from mutations at multiple sites in the genome, we used elevated mutation rates (4*N*_*e*_*µ >* 1) in the two-loci model, which allowed us to dissect the dynamics of multiple incompatible alleles segregating within populations. Under these conditions, we showed that epistatic interactions among segregating BDMi delays their fixations, especially when they are unlinked and when they are not too recessive. These results were validated by explicit multi-loci simulations. This effect was not predicted by previous approximations that showed that, when mutations are rare enough, RI only depended on mutation rate, independently on population parameters and reproductive mode [37]. However, they are in agreement with some phenomenological models that assumed that incompatibilities was function of the genetic distance between individuals, so can be counter selected in large polymorphic populations [58].

The purging of segregating BDMi mutations bears implications for the effect of selfing on RI. First, selfing increases drift and reduces polymorphism (including for BDMi mutations) and second, selfing reduces genetic shuffling thus limits the possibility for two mutations arose in different individuals and genetic background to be gathered in a same genotype. The combination of both effects facilitates RI compared to selfing, even under certain conditions where outcrossing favours the fixation of locally selected BDMi mutations.

### Mating systems and the pace of speciation

It was previously unknown if and how selfing affects the pace of speciation. Our results overall suggest that selfing reduces the fixation time of underdominant, compensatory, and BDMi mutations in allopatry, making speciation overall faster in selfing populations. Remarkably, this effect may even persist in the face of local adaptation, suggesting that – contrarily to our initial hypothesis – ecological speciation may occur faster in selfing populations than in outcrossing populations. It is unclear what are the relative importance of underdominant, compensatory, and BDMi mutation in determining the pace of speciation in natural populations. In any case, selfing broaden the spectrum of incompatibilities that can fix, and so should on average shorten the waiting time to complete speciation, even for ecological speciation.

However, the clear condition under which outcrossing should promote speciation is when it is driven by genomic or sexual conflicts, which may completely vanish under complete selfing. It is for instance known that sexual conflicts over maternal provisioning during seed development are usually stronger in outcrossers than in selfers (e.g. [59]), so that the sexually antagonistic co-evolution between male and female traits is expected to go faster in outcrossing *vs*. selfing populations, and thus to promote speciation more in outcrossing *vs*. selfing lineages [60]. A more explicit analysis remains to be done but our basic model (where selection linearly decreases with the selfing rate) confirms this prediction as soon as antagonistic selection is strong enough (say of the order of *N*_*e*_*s >* 5).

So far, empirical results are still limited but tend to support these predictions, for example with the accumulation of numerous incompatibilities between recently diverged population of selfing arctic species [18, 19] or with macro-analyses suggesting higher speciation rates in selfing lineages in Solanaceae [12, 16] as mentioned in the introduction. However, the underlying process of speciation remains unknown. In arctic species, divergence in allopatry is likely, but in Solanaceae, selfing may have promoted speciation through the limitation of gene flow, which we did not study here.

### Mating systems and the genetic architecture of reproductive isolation

Beyond the effect of mating systems on the pace of speciation, our outcomes clearly suggest that mating system should also affect the genetic architecture of speciation. In particular, underdominant and compensatory mutations are expected to be found relatively more often as reproductive barriers among selfers than among outcrossers. Classical examples of genetic modifications leading to underdominant effects include chromosomal rearrangement, which can generate and maintain RI between populations or species [28, 61]. To our knowledge, there is no studies specifically comparing the occurrence of underdominant chromosomal rearrangement in selfing *vs*. outcrossing species. Reproductive isolation due to underdominant chromosomal rearrangement is however more often found in plants than in animals [62], which is possibly due to a higher frequency of selfing plant species. Our results also suggest that reproductive barriers caused by a few strongly underdominant mutations are more likely to differ between mating systems than reproductive barriers caused by many weakly underdominant mutations.

Compensatory effects are often discussed in the context of the evolution of gene expression for which stabilising selection may lead to the co-evolution of *cis*- and *trans*-regulatory mutations (*e*.*g*., a *cis*-regulatory mutation increasing gene expression may be compensated by a *trans*-regulatory mutation decreasing gene expression, or *vice versa*) [63]. Although there compensatory mutations are expected to take a long time to get fixed [29, 40], co-evolution of *cis*- and *trans*-regulatory mutations have been found to contribute to RI between (outcrossing) species of *Drosophila* [30] *o*r mice [32], and between (selfing) species of nematode [31].

Finally, our models predict that in outcrossing species BDMi mutations are more likely to fix when they are clustered (but in repulsion) than when they are widespread. On the contrary, there is no specific constraint on genomic location in selfing species such that pairs of incompatible alleles could arise everywhere in a genome. This conclusion resembles the prediction that genes involved in local adaptation [64] or in domestication [65] should be less clustered under selfing than under outcrossing.

## Conclusions

Our analytical and simulation models show that selfing overall fosters the accumulation of underdominant, compensatory, and BDMi mutations in allopatry. This outcome help us predicting the speciation rates as well as the architecture of RI of selfing *vs*. outcrossing species. Our results bring a theoretical background to long-standing ideas [1, 5] and are tentatively supported by both phylogenetic studies and crossing experiments – though additional empirical work is needed. Future theoretical work will need to account for the effect of selfing on the stability of RI in the face of gene flow.

## Supporting information

**Fig. S1 Effects of selfing on the accumulation of underdominant mutations**.

**Fig. S2 Effects of selfing on the accumulation of a pair of compensatory mutations when the strength of the deleterious effect (***s*_*c*_**) is low (two loci model)**.

**Fig. S3 Effects of selfing on the accumulation of a pair of compensatory mutations (two loci model)**.

**Fig. S4 Effects of selfing rate on the fitness of a population, and the path taken on the fitness landscape, over the 4200 generations preceding the fixation of the pair of compensatory mutation (two loci model)**.

**Fig. S5 Effects of background selection on the accumulation of compensatory mutations (two loci model)**.

**Fig. S6 Effects of selfing on the accumulation of compensatory mutations (multi loci model)**.

**Fig. S7 Effects of selfing and selection on the accumulation of BDMi mutations (multi loci model)**.

**Fig. S8 Effects of selfing and selection on the fixation time and the number of segregating of BDMi mutations in a population (multi-loci model)**.

**Fig. S9 Effects of selfing, selection and background selection on the time to fixation fixation of BDMi mutations in a population (multi-loci model)**.

**Appendix S1 Mathematical derivations**

**File S1 Mathematica notebook**.

## Code availability

Mathematica script, C++ script, SLiM script.

## Acknowledgments

We acknowledge the GenOuest bioinformatics core facility (https://www.genouest.org) for providing the computing infrastructure. Funding was obtained from the Research Council of Norway (https://www.forskningsradet.no) to CB (project 274607: Speciation-Clock—How fast does the ‘speciation clock’ tick in selfing versus outcrossing lineages?). The funders had no role in study design, data collection and analysis, decision to publish, or preparation of the manuscript.

